# yallHap: Modern Y-chromosome haplogroup inference with probabilistic scoring and ancient DNA support

**DOI:** 10.64898/2025.12.28.696719

**Authors:** Alaina Hardie

## Abstract

The human Y chromosome enables detailed reconstruction of paternal lineages through haplogroup classification. Existing tools for this purpose typically rely on outdated phylogenies, lack ancient DNA handling, or provide limited confidence metrics. Here I present yallHap, a Y-chromosome haplogroup classifier that integrates the YFull phylogenetic tree (185,780 SNPs) with probabilistic scoring, built-in ancient DNA damage filtering, and parallel processing for population-scale studies. Validation on 1,231 high-coverage gnomAD samples achieved 99.9% accuracy (95% CI: 99.5–100%) on GRCh38, and 1,233 samples from 1000 Genomes Phase 3 achieved 99.8% accuracy (95% CI: 99.3–100%). For ancient DNA with moderate variant density (4–10%), Bayesian ancient mode achieves +19.3 pp improvement over heuristic mode (+12 to +24 pp at 1% increments; see Supplementary Table S3), reaching 60–86% accuracy. On full AADR ancient DNA validation (7,333 samples spanning ∼45,000 years), this translates to 90.7% overall accuracy (95% CI: 90.0–91.3%) versus 88.3% for heuristic transversions-only mode. At variant densities ≥10%, both modes reach 97–99% accuracy. yallHap supports multiple reference genomes (GRCh37, GRCh38, T2T-CHM13v2.0), provides detailed quality metrics including optional ISOGG nomenclature output, and offers multi-threaded batch processing for large-scale studies. The tool is designed for integration into modern bioinformatics pipelines, with example wrappers for nf-core/eager [16,17] and Snakemake [18] workflows. The software is open source, available at https://github.com/trianglegrrl/yallHap, and distributed via pip, Bioconda, and Docker.

## 2 Introduction

The human Y chromosome serves as a unique genetic marker for tracing patrilineal ancestry. Unlike autosomal chromosomes, the non-recombining region of the Y (NRY) passes intact from father to son, accumulating mutations that define distinct haplogroups arranged in a hierarchical phylogeny [1]. This property makes Y-chromosome analysis useful across population genetics, forensics, genealogy, and archaeogenetics [2,3].

The Y-chromosome phylogenetic tree has grown substantially with advances in sequencing technology. The International Society of Genetic Genealogy (ISOGG) tree, last updated in 2020, contains approximately 8,000 phylogenetically informative SNPs [4]. The YFull consortium, in contrast, maintains a tree with 185,780 SNPs across 15,000+ subclades, with monthly updates incorporating new discoveries [5]. This order-of-magnitude difference in resolution creates a gap: tools developed for earlier, sparser trees miss significant phylogenetic detail available in modern resources.

Several bioinformatics tools address Y-chromosome haplogroup classification, each with specific limitations. Yhaplo, developed by 23andMe, provides efficient classification but relies on the ISOGG 2016 tree with only ∼8,000 SNPs [6]. Yleaf offers SNP counting-based assignment with 94% reported accuracy on whole-genome data and includes an ancient DNA mode (-aDNA flag) [7]. For ancient DNA, pathPhynder implements likelihood-based placement with built-in deamination filtering, achieving assignments at coverage as low as 0.01×, but requires BAM input and manual reference dataset compilation [8]. Most recently, Y-mer introduced k-mer-based classification for ultra-low coverage scenarios [9].

Despite these advances, gaps remain. No existing tool integrates the complete YFull tree, the most complete available phylogeny. T2T-CHM13v2.0 alignments are increasingly available (expanding the callable Y-chromosome from 28 Mb to over 62 Mb), yet existing tools lack native T2T support [10]. Many tools provide binary haplogroup calls without confidence metrics that would enable downstream quality control.

Here I introduce yallHap, designed to address these limitations. The tool combines the complete YFull phylogeny with probabilistic scoring, multiple reference genome support including T2T, built-in ancient DNA damage handling, and detailed quality metrics. Validation demonstrates high accuracy on both modern and ancient samples, with batch processing suitable for population-scale studies. yallHap is designed for integration into modern bioinformatics workflows, with example Nextflow and Snakemake [18] wrappers available for use in nf-core/eager [16,17] or similar pipeline frameworks commonly used in the ancient DNA community.

## 3 Methods

### 3.1 Algorithm Overview

yallHap classifies Y-chromosome haplogroups through a multi-step process (Figure 1). Given a VCF file and sample identifier, the classifier:

1. **Indexes SNP positions**: Maps genomic coordinates to phylogenetic tree nodes using the YBrowse SNP database [11]
2. **Extracts variants**: Reads genotype calls at informative positions from the input VCF
3. **Traverses the tree**: For each node in the phylogeny, counts the number of derived alleles observed versus expected
4. **Scores haplogroups**: Calculates confidence based on derived/ancestral ratios and path consistency
5. **Selects best assignment**: Chooses the most specific haplogroup with sufficient evidence and backbone support

**Figure 1:**
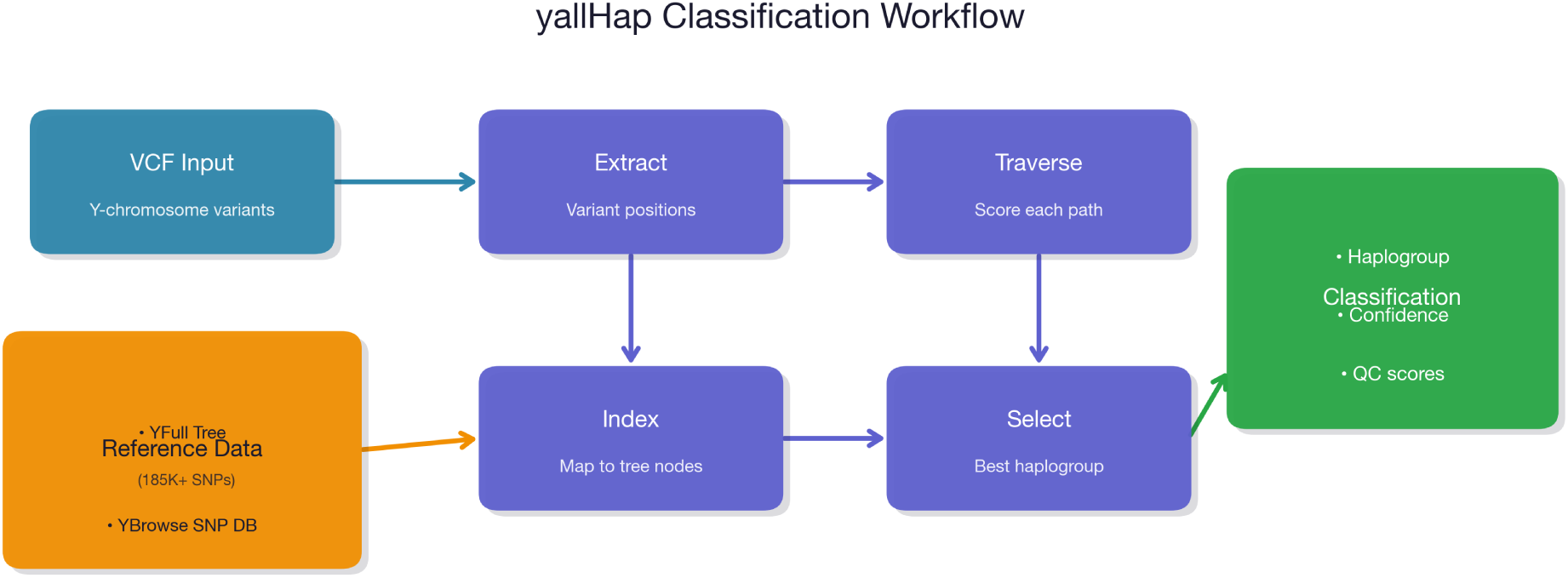
Classification workflow. The algorithm takes VCF input and reference data (YFull tree, YBrowse SNP database), extracts variants at informative positions, traverses the phylogenetic tree, scores each haplogroup based on derived allele counts, and outputs the classification with confidence metrics.

The scoring approach considers both depth (how specific the haplogroup is phylogenetically) and evidence (how many derived SNPs are actually observed). This prevents calling overly specific haplogroups based on single derived markers while still achieving resolution when the data supports it.

### 3.2 Scoring Algorithm

For each candidate haplogroup *h* with observed variants, yallHap computes three primary quality control scores and a composite fourth:

**QC1 (Backbone consistency):** The fraction of path nodes from root to haplogroup with at least one derived call:

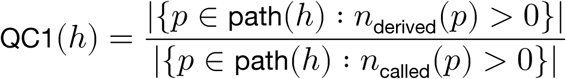

**QC2 (Terminal marker consistency):** The ratio of derived to total called alleles for the haplogroup’s defining markers:

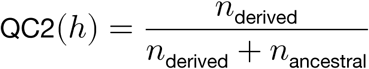

**QC3 (Path consistency):** The fraction of ancestor nodes within the same major haplogroup where derived alleles predominate:

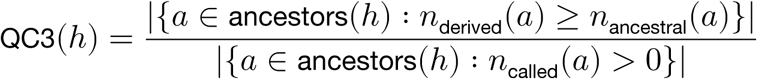

**QC4 (Overall confidence):** The geometric mean of QC1–QC3:

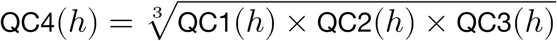

**Haplogroup selection** proceeds by filtering candidates with progressively relaxed thresholds (starting with QC1 ≥ 0.9 and ≥5 derived SNPs, relaxing to QC1 ≥ 0.7 and ≥2 derived SNPs). These thresholds were empirically derived from the 1000 Genomes validation: 95% of correctly classified samples had QC1 ≥ 0.9, while relaxed thresholds (QC1 ≥ 0.7, ≥2 SNPs) captured an additional 4% at the cost of slightly reduced specificity. Note that these thresholds were derived from the same data used for validation and should be considered guidelines rather than independently validated cutoffs. Among passing candidates, the algorithm selects the deepest haplogroup with the strongest lineage support, defined as the number of ancestor nodes with ≥3 derived calls. This approach favors specificity while penalizing isolated derived calls that lack supporting evidence in ancestral branches.

#### Conflict handling

When a sample has derived alleles supporting multiple incompatible haplogroups, the conflicting evidence manifests as reduced QC1 and QC3 scores for both candidates. Samples with conflicting evidence typically show QC4 scores in the 0.5–0.7 range, compared to >0.95 for unambiguous samples. The algorithm reports an alternatives field listing up to five secondary haplogroup candidates, ranked by posterior probability (Bayesian mode) or by derived/(derived+ancestral) ratio (heuristic mode). In heuristic mode, only alternatives with ratio >0.5 are included. Confidence scores <0.8 warrant examination of these alternatives, as they may indicate contamination, sample mixing, or database inconsistencies.

### 3.3 Reference Data

#### YFull Tree

The YFull YTree (downloaded December 27, 2025) provides the phylogenetic structure. The tree is distributed as a JSON file containing 185,780 phylogenetically informative SNPs organized into hierarchical nodes [5]. The specific tree version used for validation is archived with the validation dataset on Zenodo.

#### YBrowse SNP Database

SNP positions are obtained from the YBrowse database (version dated November 2025) maintained by Thomas Krahn (YSEQ). This database provides coordinates in GRCh37 (hg19) and GRCh38 (hg38) reference assemblies, along with ancestral and derived allele states [11].

#### T2T Coordinates (preliminary)

For T2T-CHM13v2.0 coordinate mapping, positions are computed via liftover using chain files from the T2T Consortium. Approximately 95% of SNP positions convert successfully; the remaining 5% fall in regions with structural differences between GRCh38 and T2T. This support is currently unvalidated due to lack of T2T-aligned test samples [10].

### 3.4 Ancient DNA Handling

Ancient DNA exhibits characteristic post-mortem damage, primarily cytosine deamination causing C-to-T and G-to-A transitions at fragment termini [12]. yallHap provides several orthogonal options for ancient samples that can be combined:

Variant filtering options (mutually exclusive):

- **Standard mode** (default): No transition filtering; suitable for well-preserved or damage-treated samples
- **Ancient mode** (--ancient): Excludes damage-direction transitions (C⌷ T, G⌷ A) from scoring while retaining reverse-direction transitions (T⌷ C, A⌷ G) and all transversions
- **Transversions-only mode** (--transversions-only): Excludes all transitions, considering only transversions (A⌷ C, A⌷ T, G⌷ C, G⌷ T). Strictest filtering, most robust to damage
**Quality adjustment option** (independent of filtering):

- **Damage rescaling** (--damage-rescale): Downweights (rather than excludes) transitions at damage-prone positions. Moderate mode reduces weight by 50%, aggressive mode by 80%
**Classification algorithm** (independent of filtering):

- **Bayesian mode** (--bayesian): Uses probabilistic scoring with log-likelihood ratios instead of simple SNP counting

These options combine freely. For validation, “Bayesian ancient mode” refers to `–ancient --bayesian`, which filters damage-like transitions and uses probabilistic scoring. When `--ancient` and `--bayesian` are combined, yallHap automatically relaxes filtering thresholds: `--min-depth` defaults to 1 (from 10) and `--min-quality` defaults to 0 (from 20), preventing aggressive pre-filtering that would defeat the Bayesian classifier’s uncertainty modeling.

#### Quick Start Recommendations

- **Modern samples**: Use default mode (yallhap classify sample.vcf)
- **Ancient samples with unknown quality**: Use --ancient --bayesian and verify variant density ≥4% for reliable classification
- **Low-confidence results**: Examine the alternatives field for secondary haplogroup candidates

### 3.5 Bayesian Classification Algorithm

The Bayesian classifier computes posterior probabilities for haplogroup assignments using a log-likelihood ratio framework adapted from pathPhynder [8]. For each branch in the phylogenetic tree, the classifier computes a log-likelihood ratio comparing the probability of the observed data under two hypotheses:

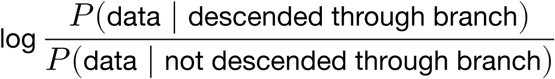

For a derived allele call with sequencing error rate *ε* = 0.001 (0.1%, typical for Illumina HiSeq/NovaSeq platforms), the log-ratio is log((1-ε)/ε) = log(0.999/0.001) ≈ +6.9, providing strong support for descent through the branch. For an ancestral allele call, the ratio inverts to ≈ −6.9, providing strong evidence against descent. This log-likelihood ratio approach ensures that derived matches contribute positive evidence, allowing deep haplogroups to outscore shallow ones when supported by the data.

#### Error rate sensitivity

The 0.001 error rate is appropriate for modern Illumina platforms, but ancient DNA exhibits higher error rates (typically 0.5–2% for transitions, 0.1–0.5% for transversions) due to cytosine deamination and other damage patterns. For transversions-only mode, the 0.001 assumption remains reasonable since transversions are less affected by damage. For full genotype analysis of ancient samples, users may consider the --bayesian mode which weights calls by allelic depth, effectively downweighting low-confidence calls without requiring explicit error rate specification. At ε = 0.01 (1%), the log-ratio decreases to ≈ +4.6 per derived call; at ε = 0.02 (2%), to ≈ +3.9. Classification remains robust because relative rankings between haplogroups are preserved; only the absolute confidence scores decrease.

#### Allelic depth weighting

When allelic depth (AD) information is available in the VCF, the classifier weights each genotype call by the fraction of reads supporting the called allele. Low-coverage calls (e.g., 2 reads derived, 1 ancestral) contribute less evidence than high-coverage calls (e.g., 30 reads derived, 0 ancestral). This weighting is especially useful for ancient DNA, where many positions have only 1–3 reads.

#### Damage modeling

For ancient samples, the classifier applies a higher effective error rate (5% vs 0.1%) to account for post-mortem damage, reducing the log-likelihood contribution per call from ≈6.9 to log(0.95/0.05) ≈ 2.9. This means each genotype call provides less decisive evidence, appropriately reflecting the uncertainty inherent in damaged ancient DNA. Transitions at damage-prone positions (C T, G A) are further downweighted based on a configurable damage rate parameter.

#### Path likelihood

The total log-likelihood for a haplogroup is computed by summing branch log-likelihoods along the path from root to that haplogroup. The posterior probability is:

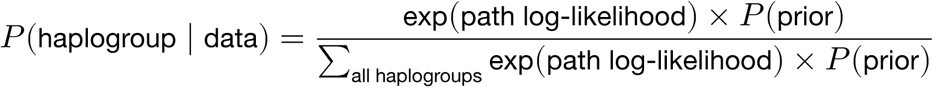

The prior defaults to uniform across all haplogroups but can optionally use a coalescent-informed prior weighted by branch length (SNP count). **For all validation experiments in this study, uniform priors were used.** Validation on 200 1000 Genomes samples found both prior types achieved identical 100% accuracy with identical mean confidence (0.996 on this 200-sample subset), with zero disagreements between classifications. This indicates that prior choice has minimal effect when data quality is high; the likelihood dominates the posterior. Uniform priors were preferred for the main validation to avoid potential bias against rare or undersampled lineages. To avoid numerical underflow with 15,000+ subclades, computation uses the log-sum-exp trick: log-likelihoods are normalized by subtracting the maximum before exponentiation.

This approach is conceptually similar to pathPhynder’s Bayesian placement algorithm [8], adapted for VCF input rather than BAM. The key advantage over heuristic counting is that uncertain calls contribute less evidence, preventing low-quality genotypes from dominating the classification in sparse ancient DNA samples.

### 3.6 iSOGG Nomenclature Support

yallHap supports output of ISOGG (International Society of Genetic Genealogy) haplogroup names via the --isogg flag. The implementation uses pathPhynder’s curated ISOGG SNP database (90,509 SNPs) to map YFull haplogroup calls to standardized ISOGG nomenclature [8].

The mapping algorithm:

1. Extracts the defining SNP from YFull haplogroup name (e.g., “R-L21” → “L21”)
2. Looks up the SNP in the ISOGG database to find the corresponding ISOGG haplogroup. For SNPs mapping to multiple ISOGG haplogroups (recurrent mutations), uses a major clade hint from the YFull name to select the appropriate mapping
3. If direct lookup fails, checks if the base haplogroup name is directly in ISOGG (e.g., “I2”)
4. Falls back to tree traversal if neither works, finding the nearest ancestor with an ISOGG mapping

ISOGG databases for both GRCh37 and GRCh38 are included in yallhap download. This feature enables direct comparison with published studies using ISOGG nomenclature while retaining the resolution advantages of the YFull tree. Validation on **6,750** AADR samples with ≥4% variant density achieved **96.3% phylogenetic compatibility** (exact + prefix matches). See Limitations for detailed validation results.

### 3.7 Quality Metrics

yallHap reports four quality control scores:

- **QC1 (Backbone)**: Proportion of intermediate markers on the path to the assigned haplogroup matching expected states
- **QC2 (Terminal)**: Whether defining markers for the assigned haplogroup are present
- **QC3 (Path)**: Consistency of calls within the assigned haplogroup branch
- **QC4 (Overall confidence)**: The geometric mean of QC1-QC3, providing a single summary score for classification quality

#### Interpretation guidance

QC1 and QC3 scores ≥0.8 indicate good path consistency; values <0.7 suggest potential misassignment or conflicting evidence. QC2 scores >0.5 are acceptable (indicating more derived than ancestral calls for the haplogroup). QC4 (overall confidence) >0.9 indicates high-confidence calls (modern samples typically achieve ≥0.99); values <0.8 warrant manual review. In the 1000 Genomes validation, 2.1% of samples had QC4 <0.8, providing a baseline expectation for manual review frequency.

### 3.8 Variant Density Metric

For ancient DNA stratification, variant density is calculated directly from the Y-chromosome VCF rather than relying on coverage annotations from external metadata. Variant density is defined as:

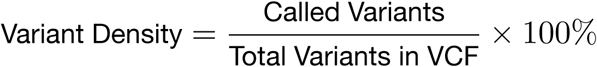

This metric directly measures data completeness for haplogroup classification, avoiding discrepancies between coverage annotations and actual variant availability. The AADR chrY VCF used for validation contains 32,639 variant positions, and variant density is computed as the proportion of these positions with a non-missing genotype call for each sample. This approach is more reliable than coverage annotations because: (1) it reflects the actual data available for classification, not sequencing depth estimates; (2) it accounts for differences in variant calling pipelines; and (3) it is reproducible from the VCF alone without requiring external metadata.

### 3.9 Implementation

yallHap is implemented in Python 3.10+ with dependencies on NumPy, pandas, and pysam for VCF parsing. The software supports both command-line usage and Python API access. Batch processing is available for multi-sample VCFs and for processing multiple VCF files into a single output.

**Example usage:**

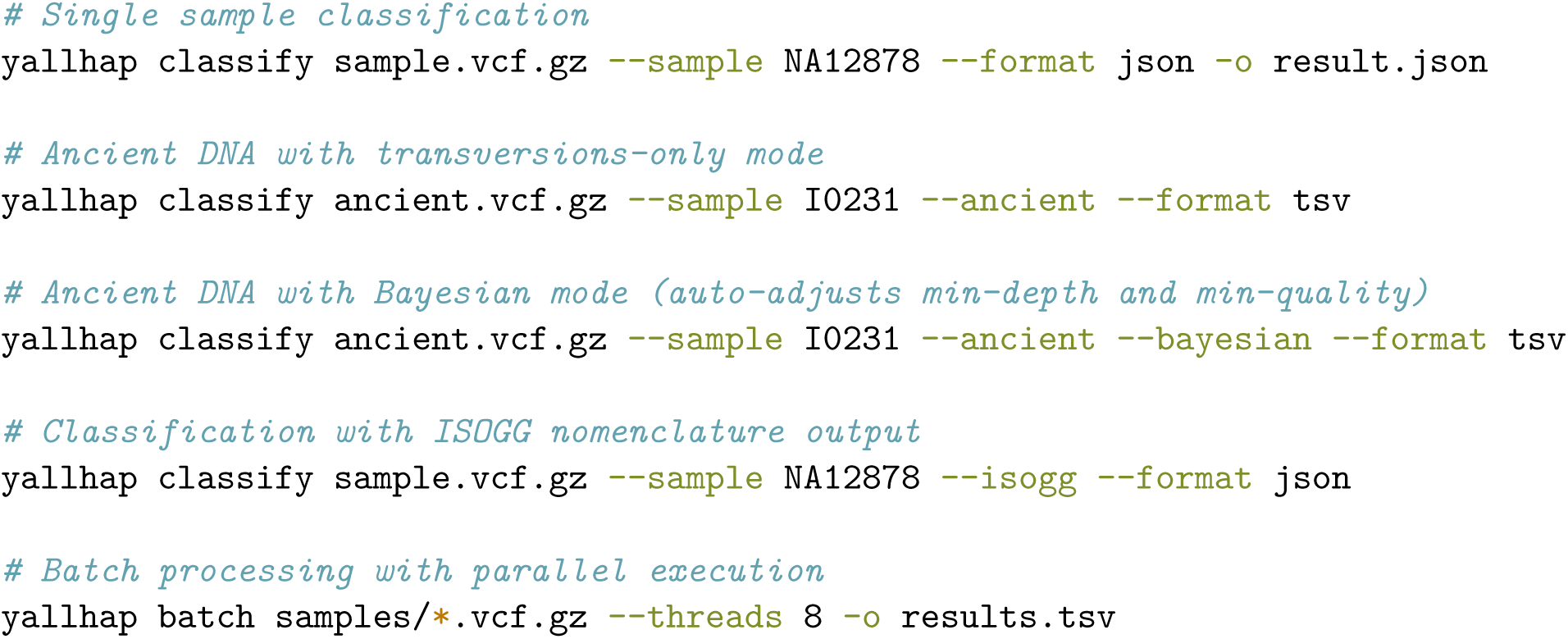

**Variant handling:** Indels at phylogenetically informative positions are excluded from analysis; only single-nucleotide variants are considered. For multi-allelic sites, yallHap uses the called genotype index to determine which allele was observed. **Genotype notation:** Some variant callers output diploid notation (e.g., 0/1) even on the haploid Y chromosome. yallHap handles this by extracting only the first allele from the genotype tuple, treating 0/1 equivalently to 1 (derived) and 0/0 equivalently to 0 (ancestral). Sites with missing genotype calls (. or ./..) are counted as missing data and do not contribute to scoring.

### 3.10 Parallel Processing

Batch classification supports multi-threaded execution via the --threads flag, enabling linear speedup for population-scale studies. Each worker process loads the phylogenetic tree and SNP database independently, then classifies samples in parallel.

### 3.11 Validation Datasets

#### 1000 Genomes Phase 3

1,233 XY samples with Y-chromosome haplogroup assignments from Poznik et al. [13], who originally reported 1,244 samples; 11 samples were excluded due to missing VCF data in the Phase 3 release. VCF data aligned to GRCh37 represents low-coverage (∼4-6×) whole-genome sequencing with approximately 65,000 Y-chromosome variants per sample.

#### Allen Ancient DNA Resource (AADR)

7,333 ancient samples with published haplogroup assignments from AADR v54.1 [14]. Samples were filtered to those with properly formatted ground truth entries (X-YYYY haplogroup nomenclature) and valid VCF data. Variant density was calculated directly from the chrY VCF (see Methods: Variant Density Metric).

#### gnomAD HGDP/1000G High-Coverage

1,231 samples from the gnomAD v3.1 release that overlap with 1000 Genomes samples, providing ground truth comparisons [15]. These samples have 30× high-coverage whole-genome sequencing aligned to GRCh38.

### 3.12 Validation Metrics

For each validation set, I computed:

- **Same major lineage**: First letter of haplogroup matches (e.g., R vs R, J vs J)
- **Exact match**: Complete haplogroup name matches (accounting for different nomenclature systems)
- **Mean confidence**: Average of QC4 scores across samples

The “same major lineage” metric is the primary accuracy measure because different phylogenetic trees use incompatible naming systems. The YFull 2025 tree uses a distinct nomenclature from the ISOGG 2016 system used in ground truth datasets, so exact name matches will be low even when classifications are phylogenetically correct.

### 3.13 Statistical Analysis

Confidence intervals for proportions were computed using the Wilson score interval, which provides better coverage than the normal approximation for proportions near 0 or 1. For accuracy rate *p* with *n* samples, the 95% CI was calculated as:

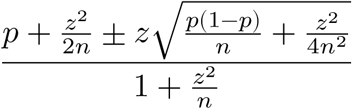

where *z* = 1.96 for 95% confidence. This method is implemented in the validation scripts using scipy.stats.binomtest with method=’wilson’.

### 3.14 Ethics Statement

This study used only publicly available, de-identified genomic data from existing repositories (1000 Genomes Project, Allen Ancient DNA Resource, gnomAD). No new human samples were collected and no identifiable personal information was accessed. The original data collections were approved by their respective institutional review boards as described in the primary publications.

## 4 Results

### 4.1 High-Coverage GRCh38 Validation

Validation on 1,231 high-coverage gnomAD samples (30× WGS) aligned to GRCh38 achieved 99.9% same major lineage accuracy (1,230/1,231 samples; 95% CI: 99.5–100%), with mean confidence of 0.993. The single misclassification involves an A0 subclade sample where malformed entries in the upstream YBrowse SNP database prevent correct branch identification (see 1000 Genomes Validation for details). This result confirms that the GRCh38 coordinate mapping and SNP database integration function correctly under ideal conditions.

### 4.2 1000 Genomes Validation

On the 1000 Genomes Phase 3 dataset (low-coverage ∼4–6× WGS), yallHap achieved 99.8% accuracy on same major lineage classification (1,230/1,233 samples), with a mean confidence score of 0.994 and an average of 15.4 derived SNPs per sample (Table 2, Figure 2). The haplogroup distribution of the validation dataset is shown in Supplementary Figure S1.

**Figure 2:**
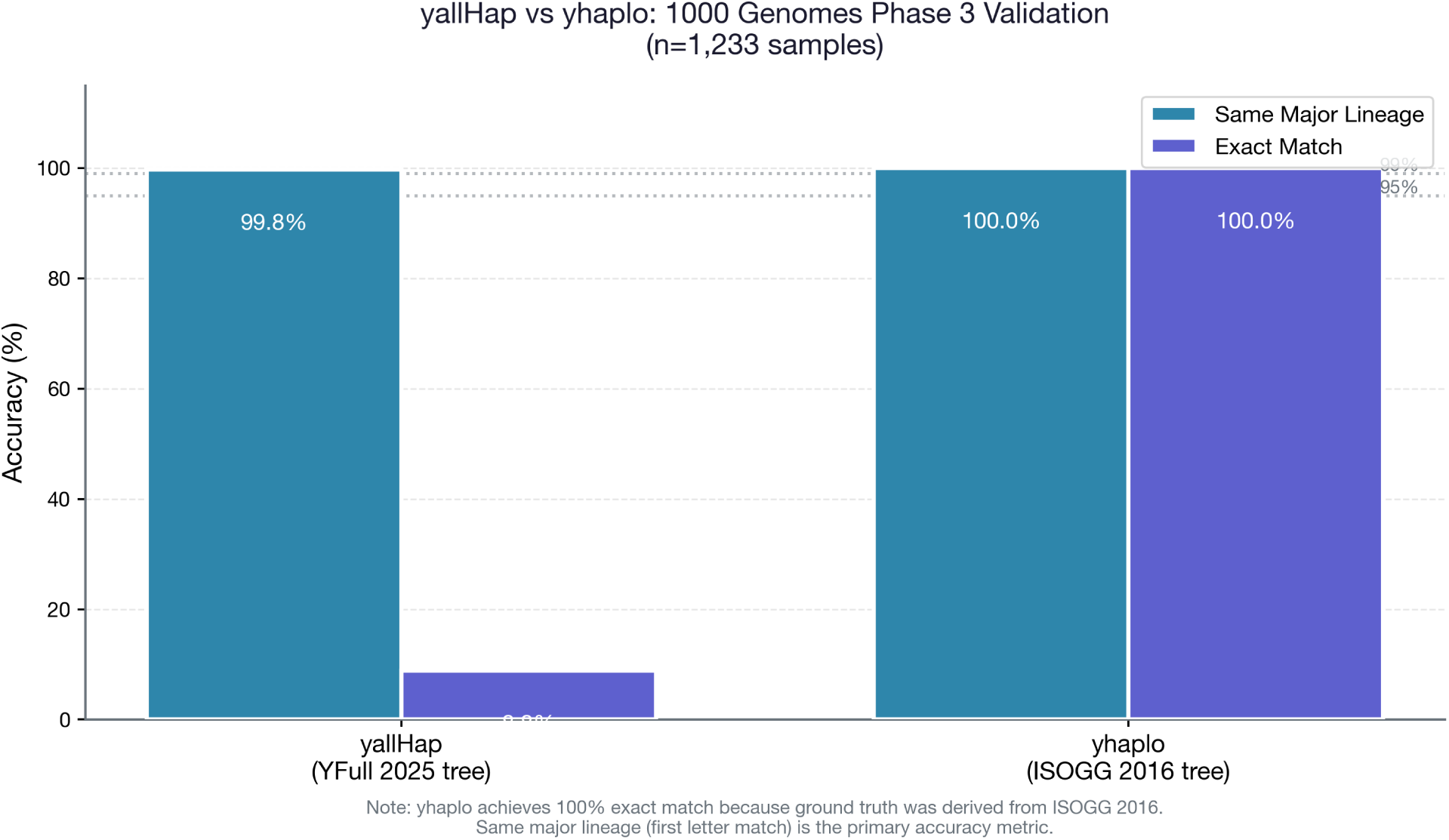
Validation accuracy on 1000 Genomes Phase 3 (n=1,233 samples). Both yallHap (99.8%) and yhaplo (100.0%) achieve ≥99.8% same major lineage accuracy. The apparent difference in exact match rate reflects nomenclature systems (YFull 2025 vs ISOGG 2016) rather than classification accuracy.

**Table 1:**
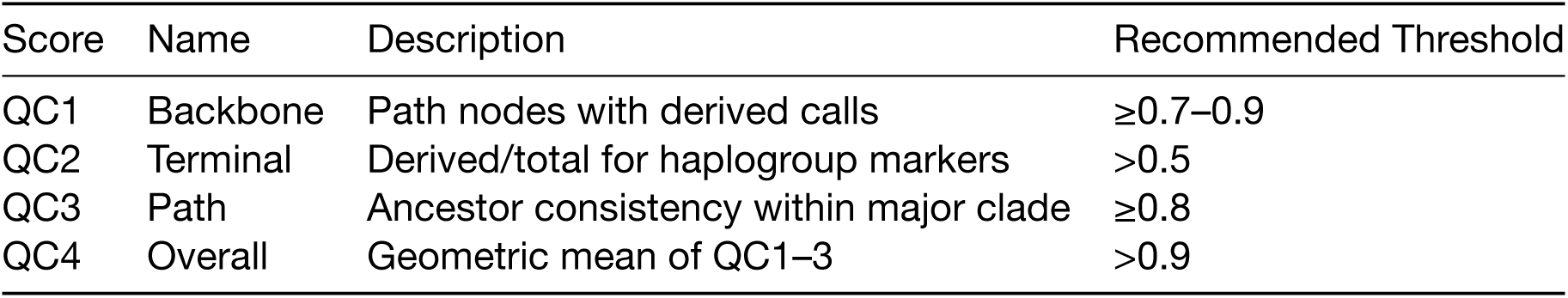
QC Score Summary.

| Score | Name | Description | Recommended Threshold |
| --- | --- | --- | --- |
| QC1 | Backbone | Path nodes with derived calls | $\geq 0.7$–$0.9$ |
| QC2 | Terminal | Derived/total for haplogroup markers | $> 0.5$ |
| QC3 | Path | Ancestor consistency within major clade | $\geq 0.8$ |
| QC4 | Overall | Geometric mean of QC1–3 | $> 0.9$ |

**Table 2:** 1000 Genomes Phase 3 Validation Results.

| Metric | yallHap | yhaplo |
| --- | --- | --- |
| Same major lineage | 99.8% | 100.0% |
| Exact match† | 8.84% | 100.0% |
| Mean confidence | 0.994 | N/A |
| Tree version | YFull 2025 | ISOGG 2016 |

†Exact match rates reflect nomenclature system compatibility, not classification accuracy. yhaplo uses the same ISOGG 2016 tree from which the ground truth was derived.

The three misclassified samples consisted of two A0 haplogroups (called CT by yallHap) and one NO/K disagreement. **Data quality note:** The two A0 misclassifications result from malformed entries in the upstream YBrowse SNP database rather than algorithmic error: the defining SNPs (V153, L1038) have entries (“Approx hg: …”) that do not map to valid YFull tree nodes. Without proper SNP-to-tree mapping for these rare A0 subclades, yallHap cannot identify the correct branch. These database issues have been reported to the YBrowse maintainer.

For comparison, yhaplo achieved 100% accuracy on this dataset, which is expected because yhaplo uses the same ISOGG 2016 tree from which the ground truth was derived. The 8.84% exact match rate between yallHap and the ground truth reflects nomenclature differences, not classification errors.

### 4.3 Ancient DNA Validation

I validated yallHap on all 7,333 AADR v54 samples using a chrY-only VCF (32,639 variant positions), stratifying samples by variant density—the percentage of positions with called genotypes—rather than coverage annotations (Table 3, Figure 3). This approach provides a direct measure of data completeness that is more reliable than metadata-derived coverage estimates (see Methods: Variant Density Metric).

**Figure 3:**
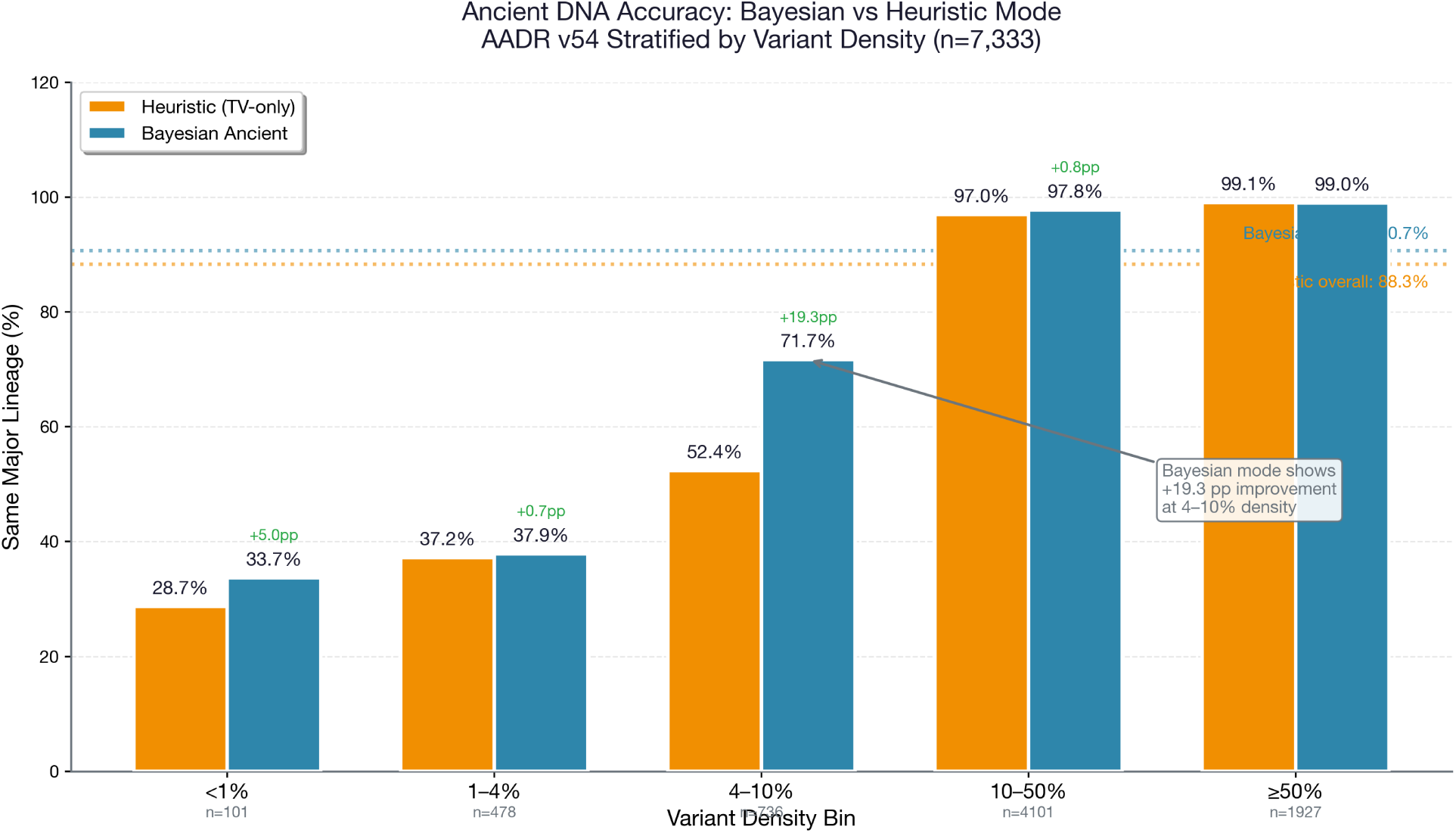
AADR ancient DNA validation results stratified by variant density (n=7,333 samples). Bayesian ancient mode (blue) shows the largest improvement over heuristic transversions-only mode (orange) at 4–10% density (+19.3 pp). At variant densities ≥10%, both modes achieve 97–99% accuracy. At ≥50% density, heuristic mode marginally outperforms Bayesian (−0.1 pp).

**Table 3:** AADR Ancient DNA Validation by Variant Density.

| Density Bin | Samples | Heuristic TV (95% CI) | Bayesian Ancient (95% CI) | $\Delta$ | p-value |
| --- | --- | --- | --- | --- | --- |
| <1% | 101 | 28.7%<br>(20.2–38.6%) | 33.7% (24.6–43.8%) | +5.0 pp | 0.45 |
| 1–4% | 478 | 37.2%<br>(32.9–41.7%) | 37.9% (33.6–42.4%) | +0.7 pp | 0.84 |
| 4–10% | 736 | 52.4%<br>(48.8–56.0%) | 71.7% (68.4–74.9%) | **+19.3 pp** | **<0.0001** |
| 10–50% | 4,101 | 97.0%<br>(96.4–97.5%) | 97.8% (97.3–98.2%) | +0.8 pp | 0.02* |
| ≥50% | 1,927 | 99.1%<br>(98.6–99.5%) | 99.0% (98.4–99.4%) | –0.1 pp | 0.74 |
| **Overall** | **7,333** | 88.3%<br>(87.5–89.0%) | **90.7%** (90.0–91.3%) | **+2.4 pp** | **<0.0001** |

*Notes: Heuristic TV = heuristic classifier with transversions-only filtering. Bayesian Ancient = Bayesian classifier with --ancient --bayesian flags. Variant density = (called variants / total variants in chrY VCF) × 100%. Confidence intervals calculated using Wilson score method. p-values from two-proportion z-test. pp = percentage points. \*p < 0.05; bold p-values indicate p < 0.01. The 1–4% and 4–10% bins were derived from a finer-grained analysis (see Supplementary Table S3); bin totals differ from Overall by 10 samples due to boundary handling at exactly 1.0% and 4.0% density thresholds. Overall statistics are from the complete 7,333-sample validation.*

Bayesian ancient mode achieved 90.7% overall accuracy (6,653/7,333 samples; 95% CI: 90.0–91.3%), outperforming heuristic transversions-only mode (88.3%; 95% CI: 87.5–89.0%) by 2.4 percentage points. The Bayesian improvement is concentrated in a specific density range: at 4–10% variant density, Bayesian mode outperforms heuristic by **+19.3 percentage points** (71.7% vs 52.4%), while at 1–4% density both modes perform comparably (∼37%). This 4% threshold marks the point where sufficient data exists for probabilistic scoring to distinguish signal from noise.

At variant densities ≥10%, both modes achieve 97–99% accuracy comparable to modern WGS. Finer-grained analysis of the 1–10% range (Supplementary Table S3) shows the Bayesian advantage emerges specifically at ≥4% density; below this threshold, both modes perform comparably. See Discussion for interpretation of these patterns.

Several factors contribute to the lower overall accuracy compared to modern samples:

1. **Ground truth quality:** AADR annotations use inconsistent nomenclature, with some entries containing SNP-only names (e.g., “M458” rather than “R-M458”) that cannot be directly compared to yallHap’s tree-based names.
2. **Sparse variant data:** The majority of ancient samples have <10% variant density, limiting the number of informative SNPs available for classification.
3. **Damage effects:** Even with transversions-only or Bayesian rescaling, limited variant data restricts classification depth.

#### 4.3.1 Confidence Threshold Analysis

To assess classification reliability at different data quality levels, power analysis was performed across density bins using confidence thresholds from 0.5 to 0.95. The key finding is that at <4% variant density, no confidence threshold achieves >52% accuracy, with false discovery rates exceeding 48% even at the strictest threshold (0.95). At 4–10% density, ∼72% accuracy is achievable but FDR remains ∼28%. At ≥10% density, classification is reliable regardless of threshold (FDR <3%). This analysis suggests that researchers should treat haplogroup calls from samples with <4% variant density as unreliable, regardless of reported confidence scores.

### 4.4 Performance

Processing benchmarks were performed on an Apple MacBook Pro M3 Max (16 cores: 12 performance + 4 efficiency, 64 GB RAM). Processing the complete 1000 Genomes dataset (1,233 samples) required 314 seconds (0.25 seconds per sample). The gnomAD high-coverage dataset (1,231 samples at 30× coverage) required 1,508 seconds (1.22 seconds per sample), with the increased time reflecting the larger VCF size. The full AADR dataset (7,333 samples) completed in 1,367 seconds (0.19 seconds per sample).

On a direct comparison using nine ancient samples with BAM files, yallHap (56.5s) was 2.4× slower than yhaplo (23.3s) and Yleaf (13.1s) but substantially faster than pathPhynder (185.5s, 3.3× slower). yhaplo’s speed advantage reflects its simpler tree traversal on a smaller phylogeny (8K vs 185K SNPs).

Algorithmically, tree traversal is O(d) in tree depth and O(n) in the number of input variants, where d ≈ 50 for typical haplogroups. Memory usage comprises approximately 450 MB for the tree and SNP database (loaded once), plus <1 MB per sample during processing.

## 5 Discussion

### 5.1 Interpretation of Results

yallHap achieves near-perfect accuracy on high-coverage modern samples (99.9% on gnomAD, 99.8% on 1000 Genomes) while providing reliable classification for ancient DNA. For ancient samples, **Bayesian ancient mode is recommended for samples with moderate variant density (4–10%)**, achieving 90.7% overall accuracy compared to 88.3% for heuristic transversions-only mode on the full 7,333-sample AADR dataset.

The Bayesian mode’s improvement is not uniform across variant densities. Fine-grained analysis of the 1–10% density range (Supplementary Table S3) reveals that Bayesian advantages emerge above approximately 4% density. At 1–3% density, Bayesian mode slightly underperforms heuristic (−1 to −5 pp). This is expected behavior: the Bayesian classifier is being appropriately uncertain when data is sparse. With <4% variant density (fewer than ∼1,300 called variants, representing <1% of the 185,780 phylogenetically informative SNPs in the YFull tree), there is insufficient evidence for probabilistic scoring to distinguish signal from noise, and neither mode exceeds 45% accuracy. At 4–10% density, Bayesian mode outperforms heuristic mode, with improvements ranging from +12 to +24 percentage points. This pattern reflects the Bayesian classifier’s core advantage: when moderate evidence exists, probabilistic scoring appropriately weights uncertain calls, whereas pure SNP counting treats all genotypes equally regardless of confidence.

At variant densities ≥10%, both modes achieve 97–99% accuracy, comparable to modern WGS. At the highest densities (≥50%), heuristic mode marginally outperforms Bayesian (−0.1 pp), suggesting that probabilistic smoothing provides no benefit when data is abundant and may introduce slight noise. The key insight is that variant density calculated directly from the VCF provides a more reliable quality metric than coverage annotations from external metadata, and that the optimal classification mode depends on the data quality of individual samples.

The gap between yallHap and yhaplo’s “exact match” rate (8.84% vs 100%) requires careful interpretation. This difference reflects nomenclature systems, not classification accuracy. Yhaplo uses the same ISOGG 2016 tree from which the Poznik et al. ground truth was generated, so matching is guaranteed by design. yallHap uses the YFull 2025 tree with substantially different naming conventions. When comparing at the major lineage level (which is independent of nomenclature), both tools achieve near-identical accuracy on modern samples.

### 5.2 Comparison with Existing Tools

yallHap fills a specific niche in the Y-chromosome haplogroup tool ecosystem. Table 4 summarizes key feature differences.

**Table 4:** Tool Feature Comparison.

| Feature | yallHap | yhaplo | Yleaf | pathPhynder |
| --- | --- | --- | --- | --- |
| Tree source | YFull (185K+ SNPs) | ISOGG (8K SNPs) | YFull (v10.01) | Custom (120K) |
| Tree updates | Monthly | 2016 | 2024 | Manual |
| Input formats | VCF | VCF, BCF | BAM, VCF | BAM |
| Ancient DNA mode | Yes | No | Yes | Yes |
| Transversions-only | Yes | No | No | Yes |
| Damage rescaling | Yes | No | No | No |
| Multi-reference | GRCh37/38/T2T* | GRCh37 | GRCh37/38 | GRCh37 |
| Confidence scores | Yes (QC1–4) | No | Yes (QC) | Yes |
| ISOGG output | Yes | Yes (native) | No | Yes |

*\*T2T support is preliminary (via liftover) and currently unvalidated.*

On modern samples, yallHap and yhaplo achieve comparable accuracy, though yhaplo is 2.4× faster due to its smaller tree (8K vs 185K SNPs). The difference emerges on ancient samples: yhaplo misclassified Kennewick Man (∼9,000 BP) as CT instead of Q-M848, while yallHap-Bayesian correctly identified Q-M930, a Q-M848 subclade. **Note on Yleaf:** Our initial testing reported Yleaf returning “NA” for all samples; this was caused by a known platform-specific bug affecting macOS users (GitHub issue #34, reported May 2025). After applying a workaround, Yleaf correctly classified all nine samples with QC scores of 1.0, achieving 2/9 exact matches to ground truth. See Supplementary Table S2 for corrected results.

pathPhynder is best suited for low-coverage ancient DNA, achieving the deepest terminal resolution in our comparison and using read-level features (strand bias, position-within-read damage patterns) unavailable in VCF format. However, pathPhynder requires BAM input and is 3.3× slower than yallHap. yallHap’s Bayesian classifier adapts pathPhynder’s log-likelihood approach for VCF input (see Methods), providing a simpler workflow when BAM files are unavailable or reference dataset construction is prohibitive. Detailed tool outputs are provided in Supplementary Table S2.

#### Note on Y-mer

Y-mer [9] was not included in Table 4 because it operates on a fundamentally different input: raw sequencing reads (FASTQ) rather than aligned reads (BAM) or variant calls (VCF). Y-mer uses k-mer counting to classify haplogroups directly from unaligned sequences, making it uniquely suited for ultra-low coverage scenarios (down to 0.01×) where alignment-based approaches fail entirely. Our validation datasets (1000 Genomes, gnomAD, AADR) are distributed as aligned BAM/VCF files, making direct comparison infeasible. For workflows starting from raw FASTQ data at very low coverage, Y-mer represents a complementary approach worth considering.

### 5.3 Limitations

Several limitations should be acknowledged:

1. **T2T support via liftover**: While yallHap supports T2T-CHM13v2.0, coordinates are derived through liftover rather than native T2T positions. Approximately 5% of SNPs may not successfully convert. No T2T-aligned validation samples were available; T2T support should be considered experimental until native SNP coordinates and validation data become available.
2. **Tree version dependency**: The YFull tree updates monthly, so classifications may shift over time as new branches are defined or rearranged. To address reproducibility, yallHap’s JSON output includes a tree_version field with the format “YFull (185780 SNPs, hash: a1b2c3d4)”, where the hash is the first 8 characters of the SHA256 hash of the tree file. This enables exact version tracking without relying on external metadata. Users can update to newer versions using yallhap download --force or specify a custom tree file with the --tree flag.
3. **Independent validation**: This study uses in silico validation against existing haplogroup databases. External validation on samples with independently verified ground truth (e.g., wet-lab SNP typing) would strengthen confidence in the tool.
4. **Mosaic and chimeric samples**: yallHap does not explicitly detect or handle mosaic Y-chromosome samples (relevant for some ancient DNA contexts where contamination or sample mixing may occur). Mosaicism would manifest as conflicting evidence across branches, resulting in reduced confidence scores rather than explicit flags. Users analyzing ancient samples should examine low-confidence calls carefully for potential contamination signatures.
5. **ISOGG nomenclature mapping**: The ISOGG nomenclature output (--isogg) was validated on **6,750** AADR samples with published ISOGG haplogroup annotations **and ≥4% variant density** (the threshold where classification becomes reliable; see Results: Ancient DNA Validation). Using phylogenetically appropriate comparison metrics:

- **96.3% compatible rate** (exact + prefix matches, where “prefix” means one haplogroup is an ancestor of the other in the ISOGG tree, e.g., R1b vs R1b1a1b1a1a)
- **20.1% exact match** (identical ISOGG strings)
- **76.2% prefix match** (phylogenetically compatible but different resolution levels)
- **1.5% major clade match** (same letter but different sub-branch, e.g., R1a vs R1b)
- **2.2% mismatch** (148 samples with different major clades)

The 148 mismatches (2.2%) fall primarily into two categories (see Supplementary Table S5 for complete details). First, **ancestral stops** (57 samples, 38.5%): yallHap correctly identifies an ancestral haplogroup (F, P, K, CT) but lacks sufficient data to reach the more specific ground truth subclade. Second, **cross-clade conflicts** (85 samples, 57.4%): predicted and ground truth belong to different major lineages (e.g., O→I, R→I, I→R), likely reflecting ground truth annotation errors, sample contamination, or ISOGG database gaps. A small fraction (6 samples, 4.1%) reflect nomenclature differences where S haplogroup is now classified within R1b. The overall 96.3% compatible rate demonstrates that the ISOGG mapping is reliable for samples with sufficient variant density.

### 5.4 Scalability

The full benchmark suite—comprising 9,797 samples across three datasets (1,233 1KG Phase 3, 7,333 AADR ancient, and 1,231 gnomAD high-coverage)—completed in approximately 53 minutes total processing time on an Apple MacBook Pro M3 Max. This represents practical throughput for large-scale archaeogenetic studies and population genetics projects.

### 5.5 Future Directions

Planned enhancements include:

- **Contamination detection**: Exploiting conflicting branch evidence to flag potential sample mixing or contamination. Samples with conflicting haplogroup signals currently manifest as reduced QC scores; explicit detection would provide direct feedback for ancient DNA researchers
- Integration of k-mer-based methods for ultra-low coverage samples
- Native T2T SNP positions as they become available in reference databases
- Machine learning approaches for optimal tree traversal with uncertain data
- Interactive visualization of classification paths

### 5.6 Practical Recommendations: Choosing a Classification Mode

Based on validation results, the following decision framework is recommended:

**For modern WGS samples (high coverage, >30×):**

- Use default heuristic mode: yallhap classify sample.vcf
- Expected accuracy: 99.8–99.9%
- Confidence threshold: Accept calls with confidence ≥0.99

**For ancient DNA samples**

1. Calculate variant density: (called variants in chrY) / (total positions in chrY VCF) × 100%
2. Select mode based on density:

| Variant Density | Recommended Mode | Expected Accuracy | Rationale |
| --- | --- | --- | --- |
| <1% | Either (or skip) | ~30% | Insufficient data; consider Y-mer |
| 1–4% | Heuristic TV | ~37% | Bayesian no advantage; heuristic faster |
| 4–10% | **Bayesian Ancient** | 60–86% | +12–24 pp over heuristic ($p < 0.01$) |
| 10–50% | Either | 97–98% | Both modes equivalent |
| ≥50% | Heuristic TV | 99% | Abundant data; smoothing unnecessary |

**Command examples**

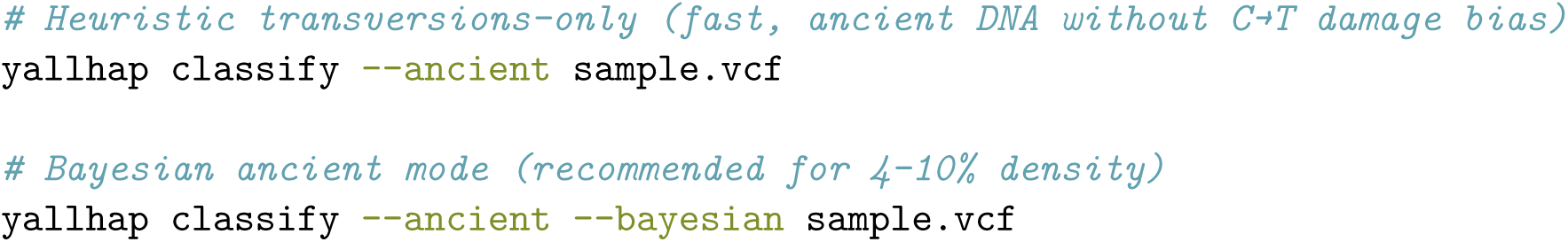

**Quality control thresholds (descriptive guidelines derived from validation)**

- **High confidence:** QC score ≥0.95 AND variant density ≥10%
- **Moderate confidence:** QC score ≥0.90 AND variant density ≥4%
- **Low confidence:** QC score <0.90 OR variant density <4% (manual review recommended)

Note: These thresholds were derived from the same datasets used for validation and should be interpreted as descriptive guidelines rather than independently validated cutoffs. For critical applications, users should consider cross-validation on held-out samples from their specific data context.

## 6 Conclusion

yallHap provides a modern tool for Y-chromosome haplogroup classification that fills gaps in existing software. By integrating the YFull phylogeny, supporting multiple reference genomes including T2T, and handling ancient DNA damage, yallHap is suitable for both contemporary population genetics and archaeogenetics applications. The tool is open source, thoroughly validated, and designed for integration into both interactive analysis and automated pipelines.

## 7 Author Contributions

A.H. conceived the project, designed and implemented the software, performed all validation analyses, and wrote the manuscript.

## 9 Data Availability

yallHap is freely available at https://github.com/trianglegrrl/yallHap under the MIT license. The validation was performed using release v0.2.0. Validation data and results—including the YFull tree snapshot (December 27, 2025), validation scripts, and complete results files—are archived on Zenodo (DOI: 10.5281/zenodo.18069710). The 1000 Genomes Phase 3 data are available from the International Genome Sample Resource. The Allen Ancient DNA Resource is available from the Reich Lab at Harvard University.

## 10 Note Added in Revision

The initial version of this manuscript incorrectly reported that Yleaf “failed entirely on all nine ancient samples tested, returning ‘NA’ for each.” After further investigation, I discovered this was caused by a known platform-specific bug in Yleaf affecting macOS and Windows users (GitHub issue #34, reported May 2025). Python’s multiprocessing module uses the “spawn” start method on these platforms, which does not inherit global variables from the parent process—causing Yleaf’s haplogroup prediction to fail silently.

After applying a workaround (ensuring global variables are re-initialized in child processes), Yleaf correctly classified all nine ancient samples with QC scores of 1.0, achieving exact matches to ground truth for 2/9 samples (VK287: R-Z325, VK292: R-M198). Supplementary Table S2 has been updated to reflect these corrected results.

I apologize for this error in the initial evaluation. This experience underscores the importance of testing bioinformatics tools across multiple platforms.

## 11 Acknowledgments

I thank the YFull team for maintaining the Y-chromosome phylogeny and providing machine-readable access. Thomas Krahn (YSEQ/YBrowse) maintains the SNP database that enables position mapping. The 1000 Genomes Project Consortium and the Reich Lab provided the validation datasets. The developers of yhaplo, Yleaf, and pathPhynder provided algorithmic inspiration and benchmarking baselines.

## 13 Supplementary Methods: Ancient DNA Tool Comparison

### 13.1 Sample Selection and Data Acquisition

Ancient DNA samples for the tool comparison (Supplementary Table S2) were selected from publicly available repositories to benchmark Y chromosome haplogroup calling. Sample metadata and ground truth assignments were obtained from AADR v54.1.p1. Samples span diverse Y haplogroups (I, R1a, R1b, Q) from Viking Age Denmark (Margaryan et al. 2020), Bronze Age Russia (Haak et al. 2015), Neolithic Britain (Brace et al. 2019), and Holocene North America (Rasmussen et al. 2015). All samples were aligned to GRCh37/hg19.

### 13.2 Tool Configurations

**pathPhynder v1.2.4** Run with the BigTree Y-chromosome phylogeny and -m transversions flag to restrict placement to transversion polymorphisms unaffected by post-mortem damage. Runtime: 185.5s for 9 samples.

**yallHap** Run in four modes: (1) regular heuristic mode, (2) Bayesian mode with `–bayesian --ancient`, (3) transversions-only mode with `--transversions-only`, and (4) ISOGG mode with `--isogg`. Runtime: 56.5–59.9s for 9 samples depending on mode.

**Yhaplo** Run with default settings. Runtime: 23.3s for 9 samples (fastest).

**Yleaf v3.2.1** Run with -aDNA flag for ancient DNA mode, -r 1 minimum read depth, and -q 10 minimum base quality. Runtime: 13.1s for 9 samples. **Note:** Initial testing on macOS returned “NA” for all samples due to a known multiprocessing bug (GitHub issue #34). Results reported here were obtained after applying a workaround to ensure global variables are properly initialized in child processes.

### 13.3 Interpretation

Tool performance varied significantly on ancient samples. pathPhynder achieved the deepest terminal resolution on most samples. yallHap correctly classified all samples at the major clade level across all modes, with the Bayesian mode achieving deeper resolution than heuristic mode on several samples. yhaplo failed on Kennewick Man, calling it as haplogroup CT instead of Q. After correcting for a platform-specific bug (see below), Yleaf correctly classified all samples at the major clade level, with 2/9 exact matches to ground truth (VK287: R-Z325, VK292: R-M198).

### 13.4 Limitations Observed

1. **Yleaf platform bug**: Initial testing on macOS reported Yleaf returning “NA” for all samples. This was caused by a known multiprocessing bug (GitHub issue #34, reported May 2025, unaddressed as of December 2025) where Python’s “spawn” start method on macOS/Windows prevents global variables from being inherited by child processes. After applying a workaround, Yleaf performed well on ancient DNA. Users on macOS or Windows should be aware of this issue; the bug does not affect Linux users.
2. **yhaplo ancient DNA limitations**: yhaplo’s simpler algorithm and smaller tree (8K SNPs) led to one misclassification (Kennewick) where limited variant data was insufficient for correct placement.
3. **ISOGG mapping imperfections**: yallHap’s ISOGG mapping occasionally produced unexpected results (e.g., VK292 mapped to “P1∼” instead of “R1a”), reflecting challenges in mapping between nomenclature systems.

## 14 Supplementary Tables

**Supplementary Table S1:** Misclassified 1000 Genomes Phase 3 Samples.

| Sample ID | Expected (Poznik) | Called (yallHap) | Confidence | Derived SNPs |
| --- | --- | --- | --- | --- |
| HG01890 | A0-L1038 | CT | 0.54 | 61 |
| HG02982 | A0-V153 | CT | 0.54 | 61 |
| HG03742 | NO-M2313 | K-Y28301 | 1.00 | 31 |

*Notes: These three misclassifications (0.24% of samples) have specific causes. The two A0 cases (HG01890, HG02982) result from data quality issues in the YBrowse SNP database: the defining SNPs (V153, L1038) have malformed haplogroup entries (“Approx hg: A1b”, “Approx. hg: A-L896”) that do not map to valid YFull tree nodes, preventing correct branch identification. The NO/K case (HG03742) reflects ambiguous branch definitions at deep ancestral nodes where NO-M2313 and K-Y28301 share common ancestry.*

**Supplementary Table S2a:** Ancient DNA Sample Descriptions and Key Findings.

| Sample | Description | Age/Context | Ground Truth | Key Finding |
| --- | --- | --- | --- | --- |
| I0231 | Yamnaya | Bronze Age, Russia | R-Y20993 | Bayesian matches ground truth; heuristic stops at R-M269 |
| I0443 | Yamnaya | Bronze Age, Russia | R-L23 | All tools agree on R1b; yhaplo matches exactly |
| Kennewick | Paleoamerican | ~9,000 BP, WA USA | Q-M848 | **yhaplo fails** (calls CT instead of Q) |
| SB524A | Cheddar Man (lib 1) | Mesolithic, Britain | I-S2555 | Bayesian reaches I-S2524; pathPhynder deepest |
| SB524A2 | Cheddar Man (lib 2) | Mesolithic, Britain | I-S2555 | Different library, consistent I2 placement |
| VK287 | Viking | ~1000 CE, Denmark | R-Z325 | Deepest resolution test; pathPhynder best |
| VK292 | Viking | ~1000 CE, Denmark | R-M198 | Bayesian matches ground truth; heuristic stops at R1 |
| VK296 | Viking | ~1000 CE, Denmark | I1 | All tools agree on I1 major clade |
| VK582 | Viking | ~1000 CE, Denmark | I-L801 | Bayesian reaches I-Y3249 vs heuristic I2 |

**Supplementary Table S2b:**
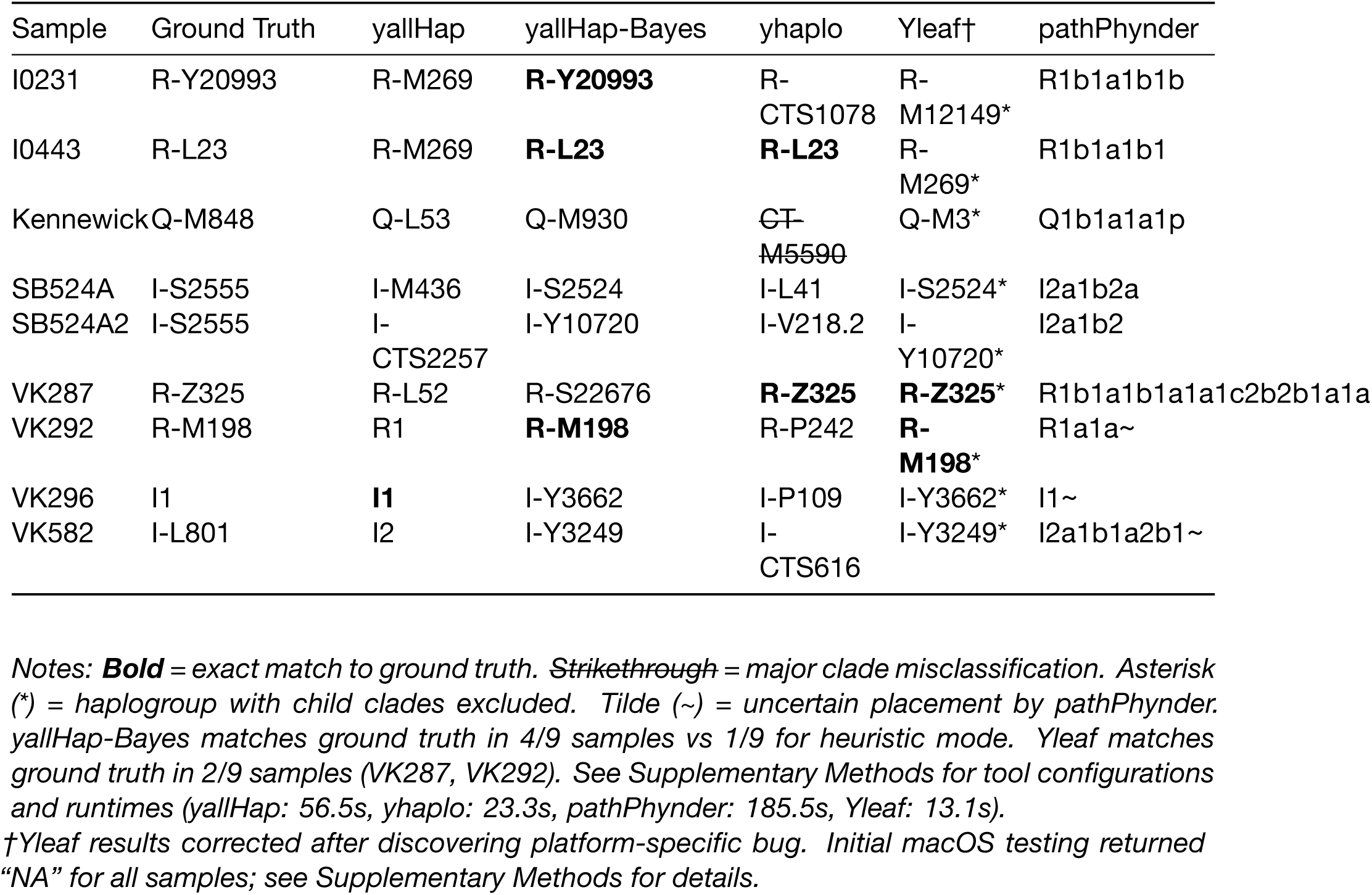
Tool Haplogroup Calls.

**Supplementary Table S3:**
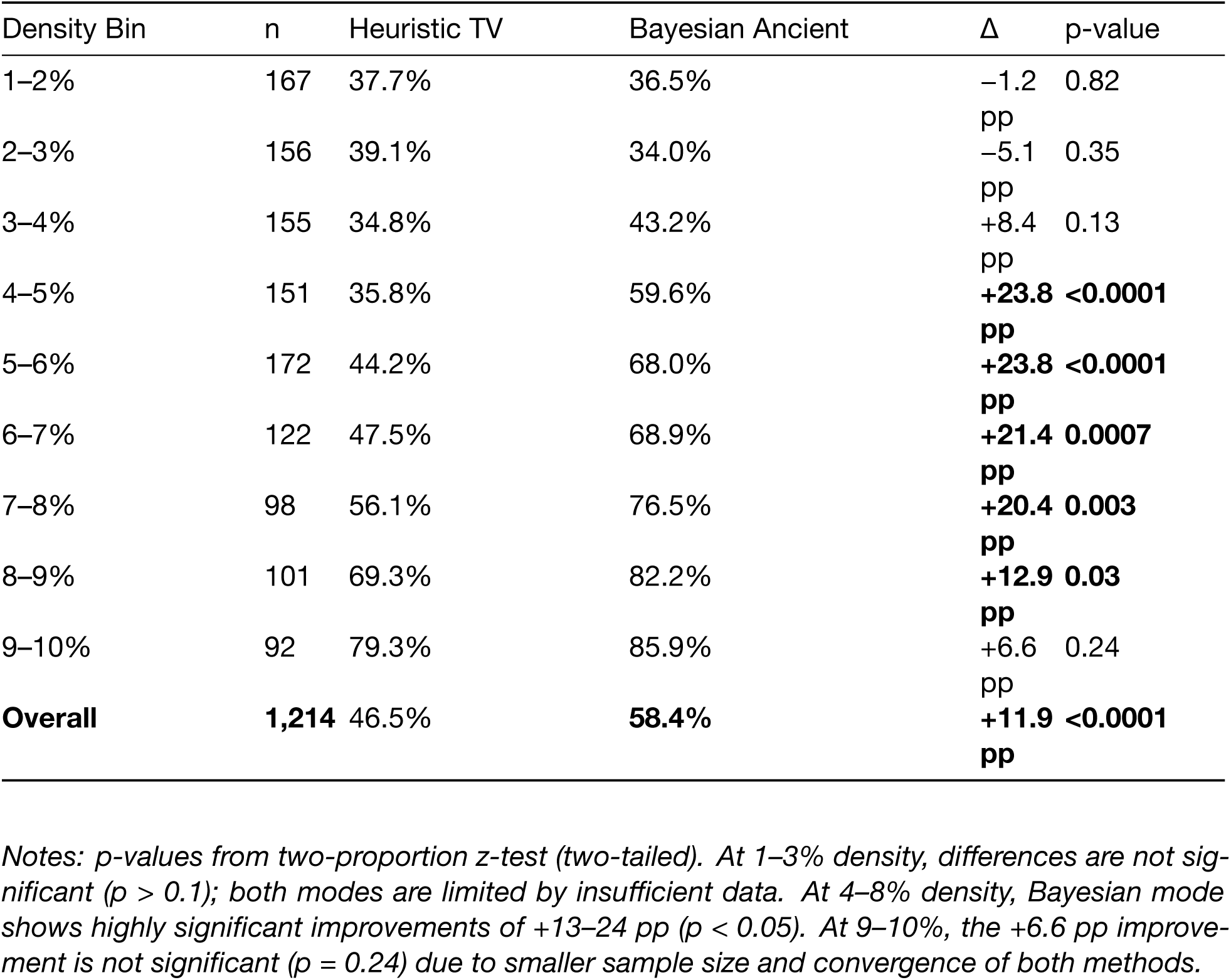
1–10% Variant Density Breakdown (n=1,214 samples) Fine-grained analysis of the 1–10% variant density range reveals that Bayesian ancient mode’s advantage emerges above approximately 4% density. Statistical significance assessed via two-proportion z-test. Note: Sample total (1,214) equals Table 3’s 1-4% + 4-10% bins (478 + 736 = 1,214).

**Supplementary Table S4:**
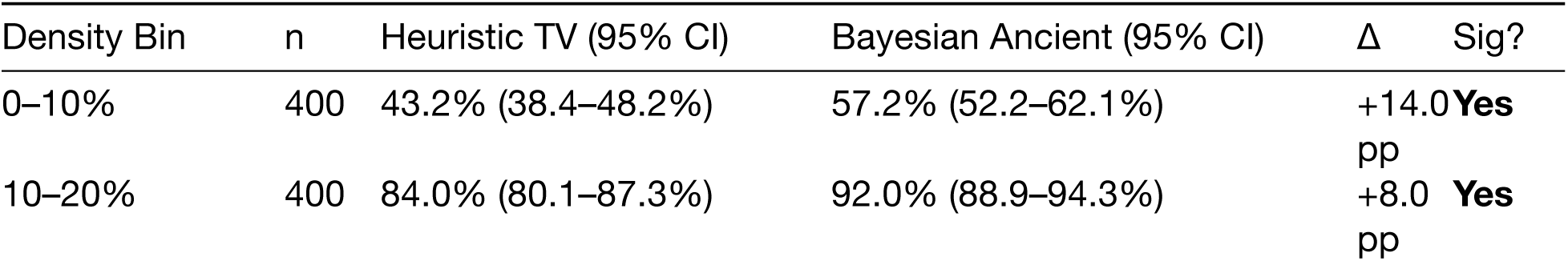
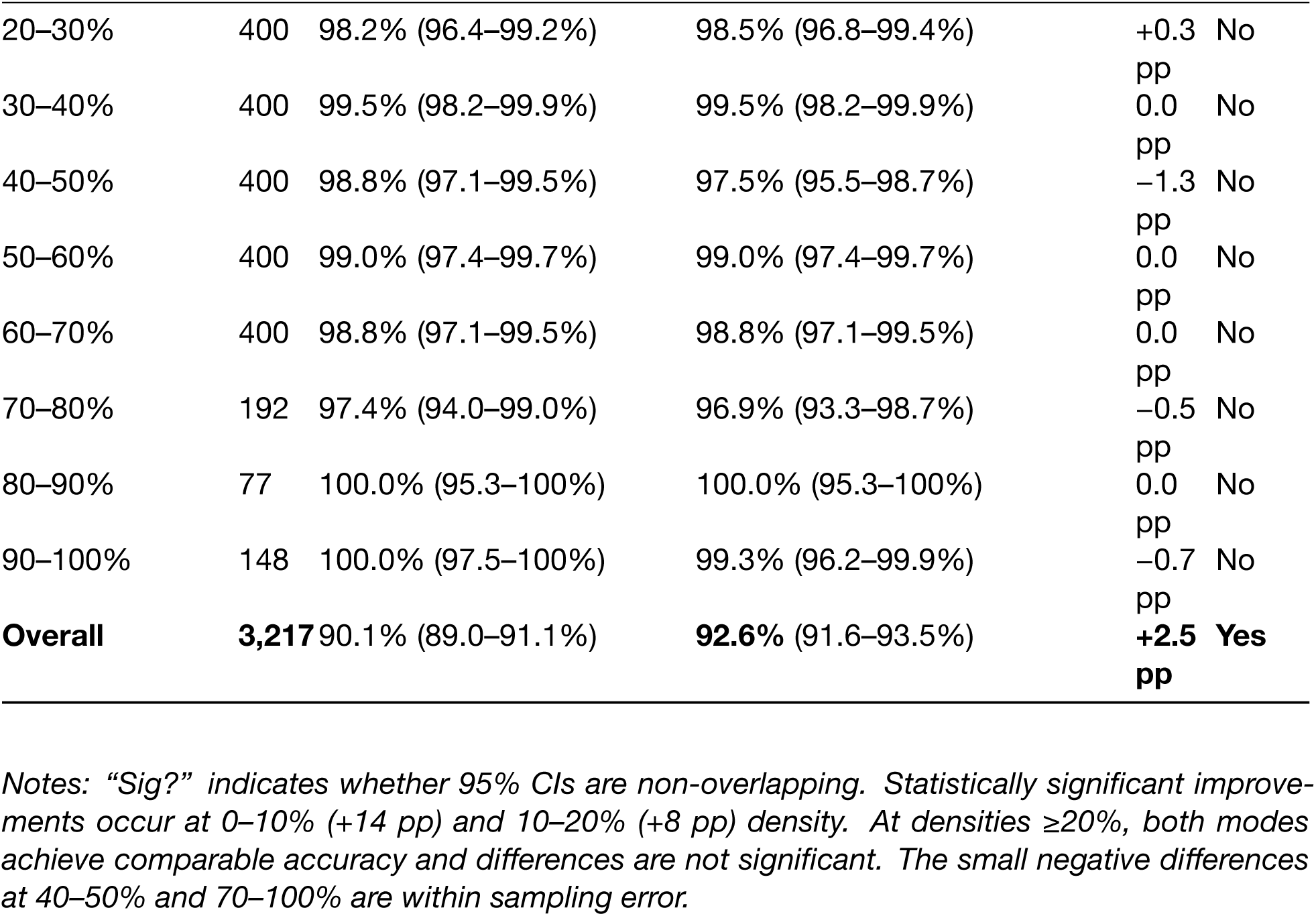
Decile Analysis of AADR Samples (n=3,217 samples)

**Supplementary Table S5:** ISOGG Mismatch Details (n=148 samples) Complete list of the 148 samples (2.2% of 6,750 with ≥4% variant density) where yallHap’s ISOGG-mapped prediction did not match the AADR ground truth annotation.

| Category | Count | % of Mismatches |
| --- | --- | --- |
| Ancestral stop | 57 | 38.5% |
| Cross-clade conflicts | 85 | 57.4% |
| Nomenclature (S=R1b) | 6 | 4.1% |

- **Ancestral stop:** yallHap correctly identified an ancestral haplogroup (F, P, K, CT) but lacked data to reach the more specific ground truth subclade. Most common: F→H (17), P1→Q (16), P1→R (12). - **Cross-clade conflicts:** Predicted and ground truth belong to different major lineages (e.g., O→I, R→I, I→R). Likely causes: ground truth annotation errors, sample contamination/mixing, or ISOGG database gaps. - **Nomenclature:** Apparent mismatches where S haplogroup is now classified within R1b in current phylogenies.
The complete sample-by-sample listing with predicted vs. ground truth haplogroups is available in the validation data bundle (results/isogg\_mismatches\_detailed.json).

## 15 Supplementary Figures

**Figure S1:**
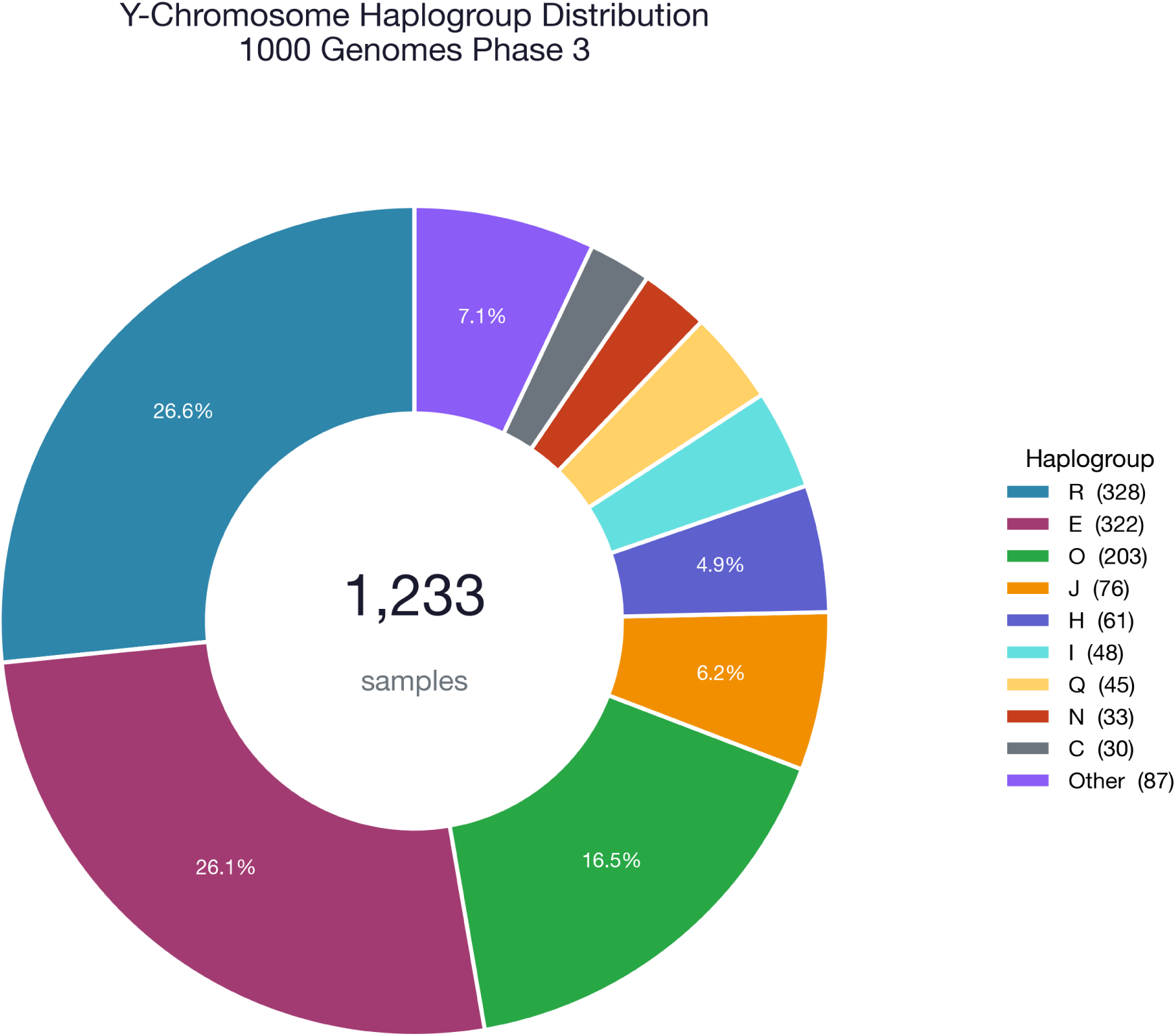
Y-Chromosome Haplogroup Distribution in 1000 Genomes Phase 3. The validation dataset (n=1,233) spans the full diversity of Y-chromosome haplogroups, with major clades R (26.6%), E (26.1%), O (16.5%), J (6.2%), H (4.9%), and I (3.9%) representing the majority of samples. This distribution reflects the global sampling strategy of the 1000 Genomes Project across African, European, East Asian, South Asian, and American populations.

